# Macroscale Thalamic Functional Organization Disturbances and Underlying Core Cytoarchitecture in Early-Onset Schizophrenia

**DOI:** 10.1101/2022.05.11.489776

**Authors:** Yun-Shuang Fan, Yong Xu, Şeyma Bayrak, James M. Shine, Bin Wan, Haoru Li, Liang Li, Siqi Yang, Yao Meng, Sofie Louise Valk, Huafu Chen

**Affiliations:** The Clinical Hospital of Chengdu Brain Science Institute, School of Life Science and Technology, University of Electronic Science and Technology of China, Chengdu, China; Otto Hahn Group Cognitive Neurogenetics, Max Planck Institute for Human Cognitive and Brain Sciences, Leipzig, Germany; Brain and Mind Center, The University of Sydney, Sydney, Australia; MOE Key Lab for Neuroinformation, High-Field Magnetic Resonance Brain Imaging Key Laboratory of Sichuan Province, University of Electronic Science and Technology of China, Chengdu, China; Institute of Neuroscience and Medicine (INM-7: Brain and Behavior), Research Centre Jülich, Jülich, Germany; Academy for Advanced Interdisciplinary Studies, Peking University; International Max Planck Research School on Neuroscience of Communication: Function, Structure, and Plasticity (IMPRS NeuroCom), Leipzig, Germany; Department of Psychiatry, First Hospital/First Clinical Medical College of Shanxi Medical University, Taiyuan, China

**Keywords:** cytoarchitectural, early-onset schizophrenia, functional hierarchy, genetic, thalamus

## Abstract

Schizophrenia is a polygenetic mental disorder with heterogeneous positive and negative symptom constellations, and is associated with abnormal cortical connectivity. The thalamus has a coordinative role in cortical function and is key to the development of the cerebral cortex. Conversely, altered functional organization of the thalamus might relate to overarching cortical disruptions in schizophrenia, anchored in development. Here, we contrasted resting-state fMRI in 99 antipsychotic-naive first-episode early-onset schizophrenia (EOS) patients and 100 typically developing controls to study whether macroscale thalamic organization is altered in EOS. Employing dimensional reduction techniques on thalamocortical functional connectome, we derived lateral-medial and anterior-posterior thalamic functional axes. We observed increased segregation of macroscale thalamic functional organization in EOS patients, which was related to altered thalamocortical interactions both in unimodal and transmodal networks. Using an *ex vivo* approximation of core-matrix cell distribution, we found that core cells particularly underlie the macroscale abnormalities in EOS patients. Moreover, the disruptions were associated with schizophrenia-related gene expression maps. Behavioral and disorder decoding analyses indicated that the macroscale hierarchy disturbances might perturb both perceptual and abstract cognitive functions and contribute to negative syndromes in schizophrenia, suggesting a unitary pathophysiological framework of schizophrenia.

## Introduction

Schizophrenia is a polygenetic psychiatric illness characterized by a combination of psychotic symptoms and motivational/cognitive deficits, which usually emerge during early adulthood (1). Over the past two decades, a wealth of neuroimaging studies has indicated that schizophrenia can be associated with pathological interactions across widely distributed brain regions, instead of focal brain damages. Accordingly, the overarching dysconnection hypothesis posits that schizophrenia results from brain structural and functional connectivity abnormalities (2). The thalamus, which is well-placed to arbitrate the interactions between distributed brain organization (3), might play a pivotal role in the pathophysiological process of schizophrenia (4, 5).

The thalamus is a cytoarchitecturally heterogeneous diencephalic structure that contains an admixture of Parvalbumin (PVALB)-rich ‘Core’ cells and Calbindin (CALB1)-rich ‘Matrix’ cells (6). Whereas core cells preferentially target granular layers (Layers III and IV) of unimodal primary regions, such as primary visual, auditory and somatosensory cortices, matrix cells target supragranular layers (Layers I-III) over wide areas in a diffuse pattern (7). This means that distinct thalamic cells may interact with cortical areas organized into different topological zones (8). Cellular-scale information of the thalamus may be a critical factor to understand the thalamocortical interactions that supports cognition and behavior. Indeed, the thalamocortical system has been suggested to form the basis for binding multiple sensory experiences into a single framework of consciousness (9). By coordinating the modular architecture of cortical networks, the thalamus has been reported to be engaged in integrating information processing within the whole cerebral cortex (10).

In accordance with its function coordinating cortical network organization, the thalamus plays the central role in the development of the cerebral cortex (11). During brain development, the thalamus changes in concordance with the cerebral cortex and disturbances of this coordinated process relate to cognitive dysfunctions (12), serving as a precursor of schizophrenia. Thalamocortical dysconnectivity patterns, characterized by hypoconnectivity with prefrontal regions and hyperconnectivity with sensorimotor areas, have been reported in both pediatric (13) and early-stage (14) patients with schizophrenia. The dysconnectivity pattern has been hypothesized to arise from disturbed brain maturation, particularly during the transition from youth to adulthood (15). Intriguingly, thalamo-prefrontal hypoconnectivity is correlated with thalamo-sensorimotor hyperconnectivity in patients, potentially implying a shared pathophysiological mechanism (14). However, few studies have investigated thalamocortical connectivity in the still developing brain of schizophrenia from a comprehensive perspective.

Recently, the application of dimension reduction techniques has emerged as a promising strategy for holistic representations of brain connectivity. These novel data-driven methods decompose high dimensional connectome into a series of low dimensional axes capturing spatial gradients of connectivity variations (16, 17). The gradient framework describes a continuous coordinate system, in contrast to clustering-based methods resulting in discrete communities (18). Using these methods in the context of cortex-wide functional connectome, previous studies observed a cortical hierarchy that spans from unimodal primary regions to transmodal regions (19), which has a close link with cortical microstructure like cytoarchitecture or myeloarchitecture (20). Their coupling along the unimodal-transmodal axis has been reported to be genetically-and phylogenetically-controlled, supporting flexible cognitive functions (21). Perturbed macroscale cortical functional hierarchies have been reported in various neurological (22, 23) and psychiatric disorders (18) including schizophrenia (24). Also, thalamic hierarchies have been previously derived from thalamocortical connectome, identifying a lateral-medial (L-M) principal gradient and an anterior-posterior (A-P) secondary gradient (25). The L-M axis captures thalamic anatomical nuclei differentiation, while the A-P axis characterizes unimodal-transmodal functional hiearchy. Also the coupling between core-matrix cytoarchitecture and functional connectome has been shown to describe the unimodal-transmodal cortical gradients, and argued to play a major role in shaping functional dynamics within the cerebral cortex (8). Given the possible implication of the thalamus in schizophrenia, thalamic hierarchies may be altered during brain maturation and could provide new insights into the disrupted thalamocortical organization in schizophrenia.

Here, we leveraged a cohort of individuals with early-onset schizophrenia (EOS), a disorder that is neurobiologically continuous with its adult counterpart (26), to examine whether macroscale thalamic functional organization shows disturbances in the still developing brain of schizophrenia, mirroring neocortical reports. To this end, we first evaluated functional hierarchies of the thalamus by employing dimension reduction techniques on thalamocortical functional connectome (27). We then embedded thalamic functional hierarchies in a neurobiological context by spatially correlating the macroscale patterns with gene expression maps from the Allen Human Brain Atlas (AHBA) (28). Last, we tested whether the functional hierarchies could estimate clinical symptoms of patients using a machine learning regression strategy.

## Materials and Methods

### Participants

Ninety-nine antipsychotic-naive first-episode EOS patients and 100 typically developing (TD) controls were recruited from the First Hospital of Shanxi Medical University and the local community through advertisements, respectively. All pediatric participants were 7–17 years old. The diagnosis of schizophrenia was in accordance with the Structured Clinical Interview for DSM-IV, and was confirmed by at least one senior psychiatrist (Y.X.) through a structured clinical interview after at least 6-months of follow-up. The psychiatric symptomatology of 71 patients was evaluated using the Positive and Negative Syndrome Scale (PANSS). Exclusion criteria for all subjects included 1) >= 18 years old; 2) neurological MRI anomalies; 3) any electronic or metal implants; or 4) substance abuse. In addition, EOS patients were excluded if they suffered from the illness for > 1 year, and TD controls were excluded if they and their first-degree relatives had any history of psychiatric disorder. This retrospective study was approved by the Ethics Committee of the First Hospital of Shanxi Medical University. Written consent was obtained from every participant and their parents or legal guardians.

Four patients and two controls were excluded from the study because of incomplete scanning data, nine patients and six controls due to excessive head motion [mean frame-wise displacement, (FD) > 0.2 mm or outliers > 50%], and one control due to poor quality of intrasubject brain registration. Ultimately, 86 EOS patients and 91 demographically-matched TD controls were included in the analysis (See **Table 1** for detailed demographic data).

**Table 1.**
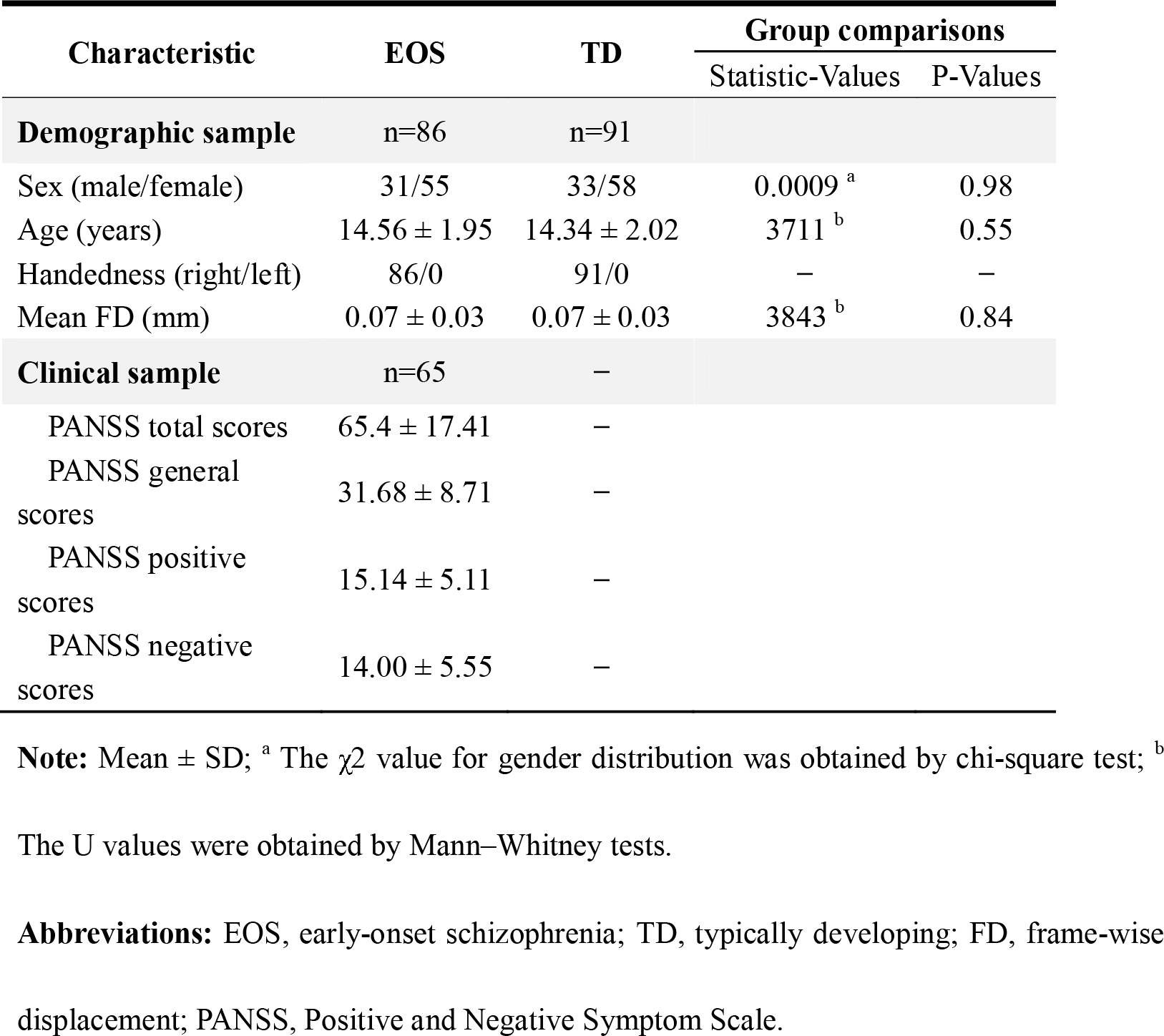
Demographics and clinical data.

| Characteristic | EOS | TD | Group comparisons |  |
| --- | --- | --- | --- | --- |
|  |  |  | Statistic-Values | P-Values |
| **Demographic sample** | n=86 | n=91 |  |  |
| Sex (male/female) | 31/55 | 33/58 | 0.0009<sup>a</sup> | 0.98 |
| Age (years) | 14.56 ± 1.95 | 14.34 ± 2.02 | 3711<sup>b</sup> | 0.55 |
| Handedness (right/left) | 86/0 | 91/0 | – | – |
| Mean FD (mm) | 0.07 ± 0.03 | 0.07 ± 0.03 | 3843<sup>b</sup> | 0.84 |
| **Clinical sample** | n=65 | – |  |  |
| PANSS total scores | 65.4 ± 17.41 | – |  |  |
| PANSS general scores | 31.68 ± 8.71 | – |  |  |
| PANSS positive scores | 15.14 ± 5.11 | – |  |  |
| PANSS negative scores | 14.00 ± 5.55 | – |  |  |

**Note:** Mean ± SD; <sup>a</sup> The $\chi^2$ value for gender distribution was obtained by chi-square test; <sup>b</sup>
The U values were obtained by Mann–Whitney tests.
**Abbreviations:** EOS, early-onset schizophrenia; TD, typically developing; FD, frame-wise displacement; PANSS, Positive and Negative Symptom Scale.

### MRI acquisition

Multimodal MRI data were collected using a 3 Tesla MRI scanner (MAGNETOM Verio; Siemens, Germany) in the First Hospital of Shanxi Medical University. T1-weighted data were acquired using a three-dimensional fast spoiled gradient-echo sequence [repetition time (TR) = 2,300 ms; echo time (TE) = 2.95 ms; flip angle = 9°; matrix = 256 × 240; slice thickness = 1.2 mm (no gap); and voxel size = 0.9375 × 0.9375 × 1.2 mm^3^, with 160 axial slices]. Resting-state functional MRI (rs-fMRI) data were obtained using a two-dimensional echo-planar imaging sequence [TR = 2,500 ms; TE = 30 ms; flip angle = 90°; matrix = 64 × 64; number of volumes = 198; slice thickness = 3 mm (1 mm gap); and voxel size = 3.75 × 3.75 × 4 mm^3^, with 32 axial slices].

### MRI processing

T1-weighted structural data were preprocessed with FreeSurfer (v7.1.0, http://surfer.nmr.mgh.harvard.edu/), which included cortical segmentation and surface reconstruction. Rs-fMRI functional data were preprocessed with the CBIG pipeline (https://github.com/ThomasYeoLab/CBIG) based on FSL [v5.0.9, (29)] and FreeSurfer (v7.1.0), which included removal of the first four volumes, slice-timing, motion correction, and boundary-based registration to structural images. See **Supplement 1** for further details. Preprocessed images were then registered to MNI152 template and resampled to the cortical surface using Ciftify package [v2.3.3, (30)]. The thalamus was localized using the Gordon 333 Atlas (31), including 2536 voxels across both hemispheres.

### Macroscale thalamocortical gradient identification

Analogous to previous work (25), gradients of thalamocortical functional connectome were generated to describe thalamic functional organization using the diffusion embedding algorithm in BrainSpace Toolbox (32). Thalamocortical functional connectivities were first calculated based on Pearson correlations between the thalamic and cortical rs-fMRI time-series for each subject (16, 23), and then converted into cosine similarity matrices (16, 17). Subsequently, nonlinear dimensionality reduction techniques were employed on similarity matrices to resolve connectome gradient, i.e., spatial axis in connectome variations (27). See **Supplement 1** for detailed analyses. We selected the first two gradients to represent the macroscale thalamic connectome space, which explained 44% of the total eigenvariance in functional connectome (**Figure S2**). The relative positioning of thalamic voxels along each organizational axis describes similarity of their functional connectivity profiles. To quantify the dispersion of each thalamic voxel in the two-dimensional gradient space, we computed eccentricity, i.e., the square root of the Euclidian distance from each thalamic voxel to the center of mass in the two-dimensional gradient space (33). For each individual, global eccentricity was calculated by averaging eccentricity values across all thalamic voxels, indicating overall dispersion of the gradient space. We additionally explored cortical-thalamic gradients by generating cortical similarity matrices from cortical-thalamic connectivity profile (see **Supplement 2** for further details).

### Thalamic functional community division

To characterize the functional relevance of macroscale thalamic gradient space, we created a thalamic functional atlas using winner-take-all representation approach (25, 34). Partial correlations were computed between rs-fMRI time-series of each thalamic voxel and six cortical functional networks (35) including the visual network (VIS), sensorimotor network (SMN), dorsal attention network (DAN), ventral attention network (VAN), frontoparietal network (FPN), and default mode network (DMN). The limbic network was excluded for low signal quality in corresponding cortical areas of our data. Each thalamic voxel was labeled by the functional network showing the highest partial correlation coefficient. Additionally, to explore cortical correspondences of thalamic gradients, we projected thalamic gradients onto the cerebral cortex. For each cortical vertex, gradient projection was calculated by correlating its cortical-thalamic connectivity profiles with thalamic gradients. These cortical maps were down sampled into 400 cortical parcels and grouped into functional networks (36) to further validate thalamic functional community division.

### The core-matrix cytoarchitecture

To delineate the core-matrix cytoarchitecture in the thalamus, we used the spatial maps of mRNA expression levels for two calcium-binding proteins (CALB1 and PVALB) (https://github.com/macshine/corematrix) (8) generated from post-mortem Allen Human Brain Atlas (28). Thalamic voxels with positive CP values (CALB1-PVALB values) related to matrix projection cells, and voxels with negative values related to core populations. Interregional correlations between CP map and differential eccentricity map were employed to reveal an association between the thalamic cytoarchitecture and disturbed functional organization in EOS patients. The variogram-matching model was used to correct for the spatial autocorrelation of brain maps (37). Subsequently, we projected CP maps onto the cerebral cortex to reveal couplings between the core-matrix cytoarchitecture and functional connectome. Cortical parcels with positive coupling values indicated as preferential associations with matrix thalamic populations, and negative values suggested core populations. To evaluate cognitive terms associated with gene-connectome coupling maps, we further conducted topic-based behavioral decoding using NeuroSynth meta-analytic database (38). See **Supplement 1** for detailed analysis steps.

### Clinical correspondences

To investigate clinical significance of thalamic functional organization disturbances, we further associated the macroscale functional phenotype with schizophrenia-related genetic expression. Based on previous work (39), we selected out 28 protein-coding genes particularly implicated in schizophrenia etiology or treatment (See **Table S1** for detailed information). Within an overlapping thalamic mask (1969 voxels), estimated mRNA levels were extracted from Allen Human Brain Atlas and then normalized using a robust sigmoid function (40). Gene expression maps were then spatially correlated with differential eccentricity map between the EOS and TD groups, corrected by the variogram-matching model (1000 surrogate maps) as abovementioned.

Second, we estimated the relationship between thalamic functional hierarchies and clinical presentations in EOS patients. Following a machine learning pipeline, we used the elastic net regression model to predict clinical symptoms in EOS (41). Eccentricity values of the two-dimensional gradient space were defined as input features, and PANSS positive and negative scores were used as predictors. The model performance was evaluated by comparison of observed and predicted clinical scores. See **Supplement 1** for detailed analyses.

### Group comparisons between the EOS and TD groups

Between-group differences on all kinds of measurements were assessed by using two-sample t-tests with covariates including age, gender, and mean FD. The multiple comparison corrections were conducted using three methods for different spatial scales: voxel-wise, parcel-wise, and global-wise. For voxel-wise thalamic gradient and eccentricity values, multiple comparison corrections were employed using the permutation-based threshold-free cluster enhancement (TFCE, 10,000 permutations, *p* < 0.005) method (FSL-PALM), which could improve sensitivity and interpretable than cluster-based thresholding method (42). For parcel-wise gene-connectome coupling values, false discovery rate (FDR) corrections (*p* < 0.005) were used to control the effect of false positives. For global-wise index like network-level gradient values and global eccentricity values, Bonferroni corrections were conducted with a significant level of *p* < 0.05.

## Results

### Macroscale thalamic gradients in TD and EOS

The principal gradient (G1, 24.6% explained) of the thalamus revealed a L-M axis, and the second gradient (G2, 11.3% explained) described an A-P axis, in line with previous work (25). In **Figure S2**, we also showed the third gradient pattern running in ventral-dorsal direction (9% explained). In EOS patients, we observed expansions at both anchors of the G1; the lateral portions including the ventral lateral and ventral posterior thalamic nuclei, the medial dorsal areas compared to TD controls (**Figure 1A**, TFCE, *p* < 0.005/2). We also observed expansions along the G2 axis including the pulvinar and anterior nuclear groups (**Figure 1B**, TFCE, *p* < 0.005/2). Combing G1 and G2 axes, we computed an eccentricity score using the square root of the Euclidian distance from each thalamic voxel to the center of mass in the two-dimensional gradient space (33). Global eccentricity was assessed for each participant by averaging eccentricity values across all thalamic voxels. We found significantly increased global eccentricity for thalamic voxels in EOS compared to TD (t = 2.34, *p* = 0.02), indicating a segregation of macroscale thalamic functional organization in patients (**Figure 1C**). Additionally, we explored the spatial pattern of cortical-thalamic gradients, reflecting the organization of cortical-thalamic connectivity in the cortex (**Figure S3**). The first cortical-thalamic gradient was similar with previous cortical functional gradients, showing a unimodal-transmodal transition pattern. See **Supplement 2** for further discussions.

**Figure 1.**
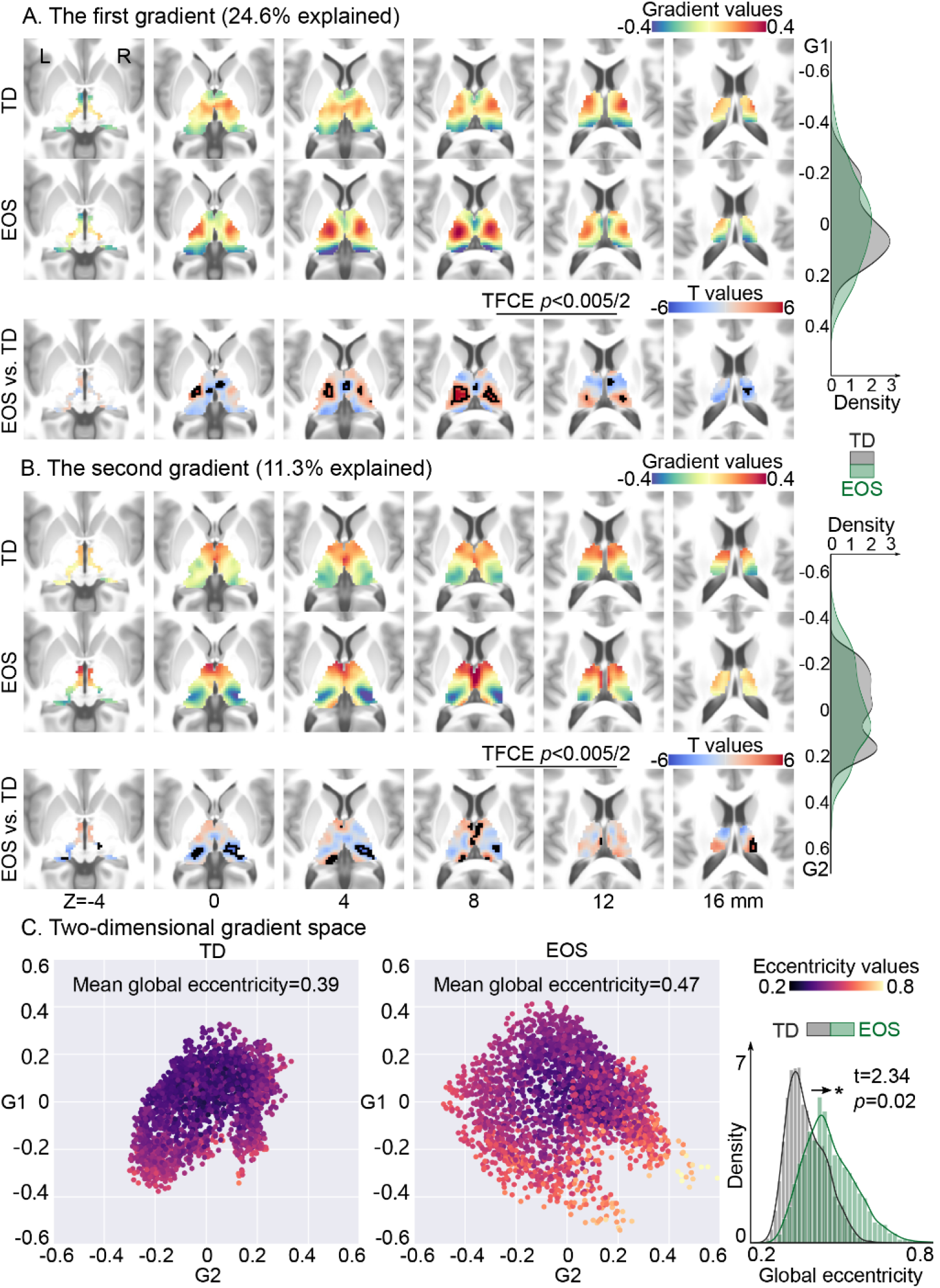
Thalamic gradients in typically developing (TD) controls and early-onset schizophrenia (EOS) patients. (**A)** The group-level primary gradients (G1) for TD and EOS, and their between-group differences. The G1 depicts a transition from lateral to medial portions of the thalamus. Thalamic voxels showing significant G1 score differences were surrounded by black contours [t-test, EOS vs. TD; threshold-free cluster enhancement (TFCE), *p* < 0.005]. The density map represents the distribution of G1 loading for EOS (green) or TD (gray). (**B)** The group-level secondary gradients (G2) for TD and EOS, and their differences. G2 separates the anterior thalamic portions from the posterior portions. Thalamic voxels with significant G2 differences were surrounded by black contours. The density map represents the G2 loading for EOS (green) or TD (gray). (**C)** Gradient spaces built on the group-level G1 and G2, separately for TD and EOS. Each point represents a thalamic voxel embedded in the gradient space. Voxels are color coded based on their mean eccentricity scores across subjects. Eccentricity score was computed by the square root of the Euclidian distance from each thalamic voxel to the center of mass in the two-dimensional gradient space. Higher eccentricity indicates greater segregation, e.g., larger dissimilarity of thalamocortical connectivity, in the gradient space. The density plot depicts the distribution of eccentricity scores in EOS and TD.

### Functional relevance

Utilizing a whole-brain functional network parcellation (35), thalamocortical functional connectome was distributed into six functional communities (**Figure 2A**). We then averaged gradient loadings within each community for each subject and compared them between the EOS and TD groups (**Figure 2B**, Bonferroni correction, *p* < 0.05/6). EOS patients had significantly higher mean G1 scores in the SMN (t = 5.37, *p* < 0.0001) and DAN (t = 2.94, *p* = 0.004), and lower G1 scores in the VAN (t = −2.85, *p* = 0.005), FPN (t = −3.69, *p* = 0.0003), and DMN (t = −3.03, *p* = 0.003) relative to TD controls. For the G2 axis, patients showed decreased gradient scores in the VIS (t = −4.08, *p* < 0.0001), and increased in the DMN (t = 5.45, *p* < 0.0001). In the two-dimensional gradient space showing functional relevance (**Figure 2C**), EOS patients had apparent dissociation of SMN-related thalamic voxels along the G1 axis, and a larger dissociation of VIS- and DMN-related regions along the G2 axis.

**Figure 2.**
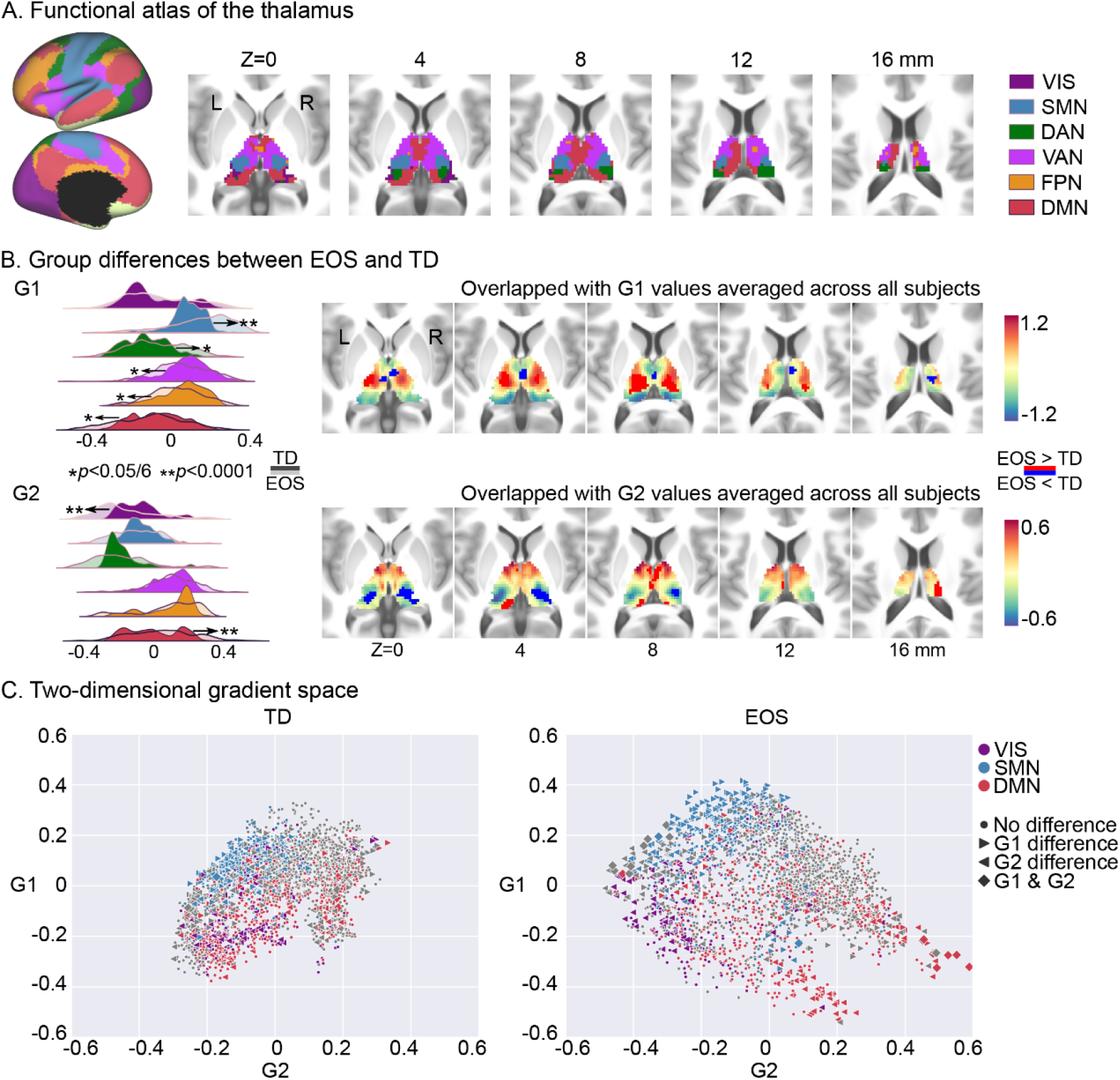
Thalamic gradients distributed into functional communities. **(A)** Network-level representations of the thalamocortical connectome. (**B)** G1 and G2 scores within functional communities. Density maps indicate gradient scores (top left: G1, bottom left: G2) within six networks for EOS and TD. Network level differences between EOS and TD were accessed by t-tests, and significant differences are depicted by * (Bonferroni correction, *p* < 0.05) and ** (*p* < 0.0001). Thalamic G1 (top right) and G2 (bottom right) representations across all participants. Voxels showing higher gradient values in EOS are colored red, whereas voxels with lower gradient values are colored blue. (**C)** Gradient space representation of the thalamus together with functional communities for TD (left) and EOS (right). Thalamic voxels are situated based on their G1 (x-axis) and G2 (y-axis) scores, and are colored according to their network assignment (purple-VIS; blue-SMN; red-DMN; grey-all other networks). ► stands for the G1 score differences between EOS and TD; ◄ for the G2 score differences; ♦ for both G1 and G2 score differences; ● for no difference. VIS, the visual network; SMN, the sensorimotor network; DAN, the dorsal attention network; VAN, the ventral attention network; FPN, the frontoparietal network; DMN, the default mode network.

Moreover, rather than dividing the thalamus into functional communities, we also projected thalamic gradients onto the cerebral cortex and evaluated their correspondence with cortical functional networks (**FigureS4**). Both cortical projections of G1 and G2 tended to follow the unimodal-transmodal transition. However, the VAN was located in the unimodal part of G1 projection, while the transmodal portion of G2 projection. Nevertheless, the dissociation of SMN-related thalamic voxels in EOS was supported by results of G1 projection, and the VIS-associated dissociation of G2 axis was also replicated.

### The core-matrix cytoarchitectural basis

Having established macroscale thalamic connectome gradients, we further investigated whether connectome differences between patients and controls were specific to thalamic cytoarchitectural features, namely core vs. matrix cells, motivated by a previous work (8). Thus, mRNA expression levels for CALB1 (core type cells) and PVALB (matrix type cells) were separately assessed at the thalamic voxel level (**Figure S5A**). The differential expression level between CALB1 and PVALB, i.e., CP scores, was used to delineate thalamic core/matrix type cell distribution, where positive/negative CP value indicated with matrix/core cell type, respectively. Eccentricity was used to evaluate the gradient space-embedded position of each thalamic voxel (**Figure S5B**). EOS patients showed increased eccentricity in medial dorsal, ventral lateral, and ventral anterior nucleus compare to TD controls, indicating their increased dispersion from the center of the gradient space (TFCE, *p* < 0.005).

We then mapped the core-matrix cytoarchitectural features onto the two-dimensional gradient space (**Figure 3A**). Core cells showed mildly increased global eccentricity in EOS patients compared with core cells of TD (t = 2.38, *p* = 0.02), but matrix cells did not (t = 1.62, *p* = 0.11). No significant difference was found between global eccentricity of core and matrix populations within EOS (t = 1.07, *p* = 0.29) or TD (t = 0.36, *p* = 0.72). Next, Pearson’s correlations were used to quantify group-level associations between the cytoarchitecture and the connectome gradient disturbances, and the variogram methods that control for the spatial autocorrelations were used to determine statistical significance levels (**Figure 3B**). A negative correlation was found between CP values and t values of eccentricity (r = −0.29, *p_vario_* = 0.002). In detail, increased dispersion of macroscale connectome gradient space in patients was particularly related to core cells (r = 0.31, *p_vario_* < 0.0001), rather than matrix cells (r = 0.02, *p_vario_* = 0.84).

**Figure 3.**
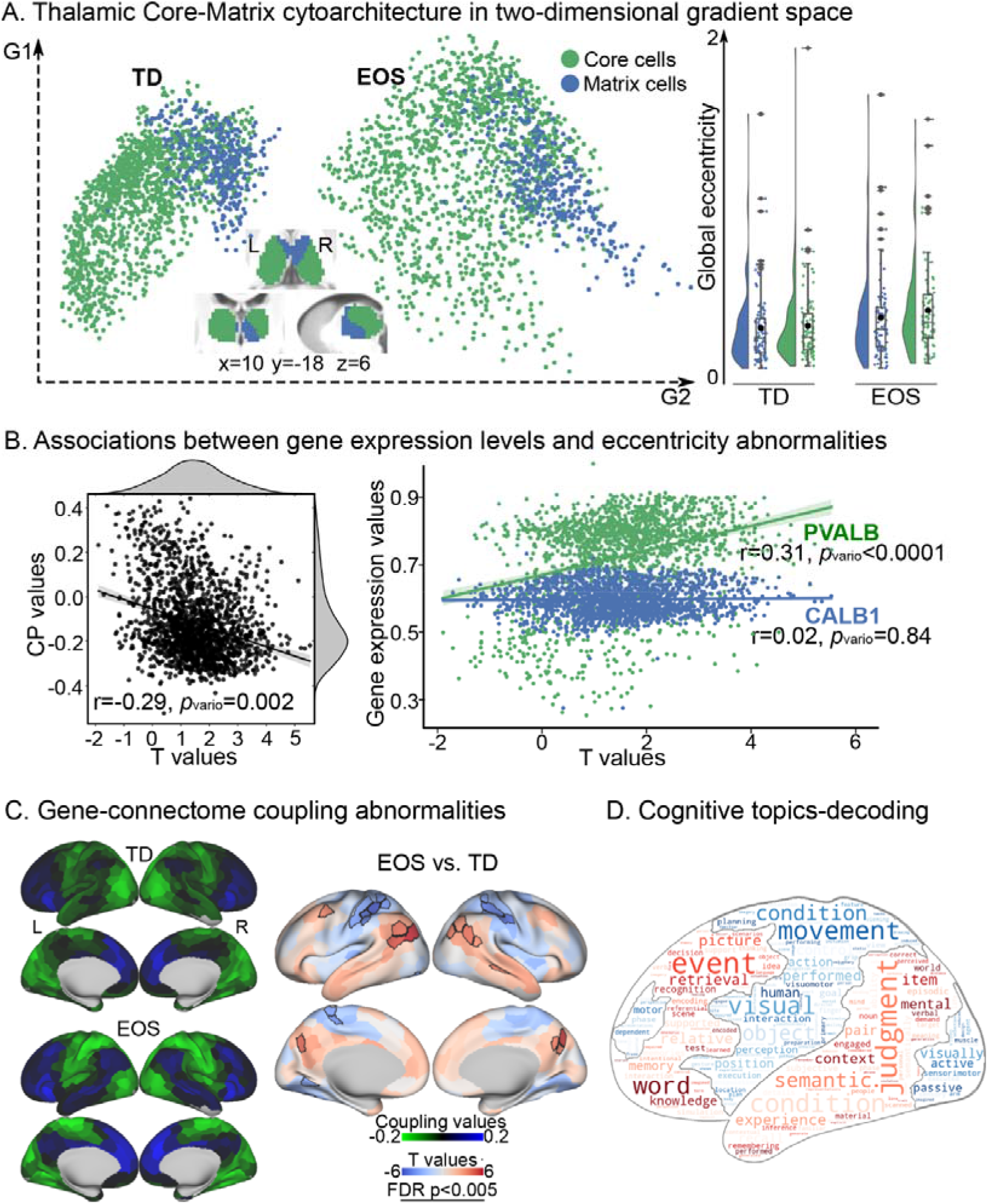
The core-matrix cytoarchitecture of the thalamus. **(A)** Core-matrix cytoarchitectural features of thalamic voxels projected onto the two-dimensional gradient space for TD and EOS. Rainclouds present group comparisons of global eccentricity in four fashions: global eccentricity values across core cells in EOS vs. global eccentricity values across matrix cells in EOS; core cells in TD vs. matrix cells in TD; core cells in EOS vs. core cells in TD; matrix cells in EOS vs. matrix cells in TD. **(B)** Associations between eccentricity differences (t values; EOS vs. TD) and differential gene expression levels (CP, left) as well as single gene expression levels (CALB1 and PVALB, right). Correlations were obtained across thalamic voxels (Pearson r values) and their significances were tested using variogram approach (*p_vario_* values). **(C)** Parcel-wise couplings between functional connectome and gene expression for TD and EOS, and their differences (t-test, EOS vs. TD). Cortical parcels with positive coupling values (blue) indicate as preferential associations with matrix thalamic cells, and negative coupling values (green) suggested core cells. Parcels with significantly different coupling patterns in patients are surrounded by black contours [false discovery rate (FDR) corrections, *p* < 0.005]. **(D)** Topic-based behavioral decoding of regions with abnormal couplings in EOS. Cognitive terms in warm color correspond to brain regions showing hyper-couplings in patients relative to controls, and cool color represent hypo-couplings. In the word cloud, the size of a cognitive term is proportional to its loading strength for decoding an input brain mask.

### Cognitive relevance

Given the association between CP map and thalamocortical connectome, we further assessed whether gene-connectome coupling contributes to cognitive processing. Thus, we projected the core-matrix cytoarchitecture to the cerebral cortex and then conducted a behavioral decoding using the NeuroSynth database (38). Couplings between functional connectome and gene expression were computed and further down sampled into 400 cortical parcels. As shown in **Figure S6**, core cell populations were mainly associated with unimodal regions that subserve primary sensory and multisensory functions. Matrix cell populations were associated with transmodal cortices characterized by more abstract cognition.

Compared with TD controls, EOS patients showed increased gene-connectome couplings in the DMN including the inferior parietal lobule, posterior cingulate cortex, inferior prefrontal cortex, temporal areas, and the FPN including the precuneus and inferior parietal sulcus, and temporoparietal junction network (**Figure 3C,** FDR, *p* < 0.005). These abnormal coupling increases were associated with seven cognitive topics including declarative memory, autobiographical memory, working memory, verbal semantics, social cognition, language, and visuospatial (z-statistic > 3.1), indicating its implications in higher-level cognitive processes. Additionally, reduced gene-connectome couplings were observed in the VIS including the extrastriate and inferior extratriate, and the DAN including postcentral regions and superior parietal lobule, as well as the SMN. These reductions were characterized by low-level visual sensory and motor functions involving visual perception, visual attention, motor, action, eye movements, visuospatial, multisensory processing, and reading (**Figure 3D**).

### Clinical relevance

To reveal clinical significance of the macroscale functional gradients, we first explored its associations with schizophrenia-related gene expression maps, based on a previous work (39) (**Figure 4A**). In particular, spatial correlations were conducted to quantify their relationships, significance levels of which were evaluated by the variogram methods. The difference map of eccentricity scores between patients and controls was significantly correlated with mRNA expression levels for Glutamatergic neurotransmission-related genes including GRIN2A (r=0.34, *p_vario_* =0.001), GRIA1 (r=-0.23, *p_vario_* =0.03), Calcium signaling-related genes RIMS1 (r=0.26, = *p_vario_* 0.005), synaptic function and plasticity-related genes including CNTN4 (r=0.32, *p_vario_* =0.001) and SNAP91 (r=0.24, *p_vario_* =0.02), other neuronal ion channels-related genes including HCN1 (r=0.25, *p_vario_* =0.03) and CHRNA5 (r=0.33, *p_vario_* =0.001), neurodevelopment-related genes including BCL11B (r=-0.23, *p_vario_* =0.02) and FAM5B(r=0.27, *p_vario_* =0.006). No significant correlation was observed in the other 19 genes, including therapeutic target-related genes.

**Figure 4.**
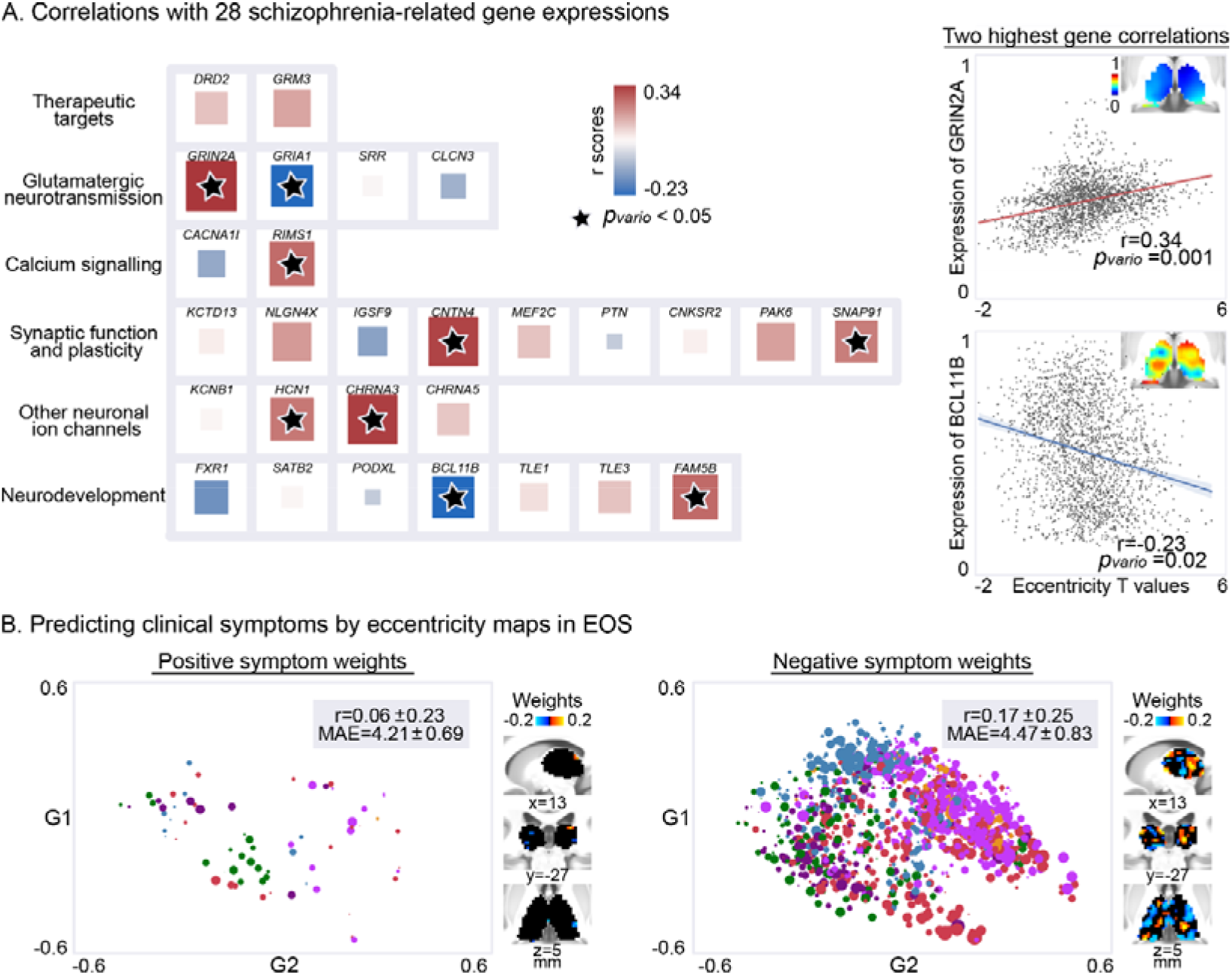
Clinical relevance of thalamic gradients in EOS. **(A)** The relationship between the macroscale functional phenotype and 28 schizophrenia-related gene expressions. Interregional correlations (Pearson r coefficients) were employed between eccentricity differences (t values; EOS vs. TD) and gene expression levels, and their significances were evaluated by variogram approach (✶ represents *p_vario_* value < 0.05). In the left panel, the size of a square is in proportion to absolute value of corresponding Pearson r coefficient, and its color is coded by the sign of r value (red-positive; blue-negative). The highest positive correlation with eccentricity differences was found at a glutamatergic neurotransmission protein-coding gene, i.e., GRIN2A, whose expression pattern is shown in the top corner. The highest negative correlation was found at BCL11B, a neurodevelopment-related gene. **(B**) The relationship between thalamic functional organization and clinical presentations. Eccentricity values in patients were used to predict PANSS positive and negative scores based on a linear regression model. The model performance was evaluated by comparing observed and predicted clinical scores. This procedure including model learning and testing was repeated 101 times, generating the distributions of Pearson r coefficients and mean absolute error (MAE). The model with median performance was reported here, and absolute value of feature coefficient was used as weight for each thalamic voxel (a point in the two-dimensional gradient space). The size of a point is coded by predictive weight, and is colored according to its network assignment (purple-VIS; blue-SMN; green-DAN; rose-VAN; orange-FPN; red-DMN).

Second, we investigated whether thalamic gradients could predict clinical symptoms in EOS. We used eccentricity scores as input features in a linear regression model to estimate PANSS positive and negative scores of patients. The optimal model parameter (L1 ratio) was 0.3 for positive symptoms prediction, and 0.1 for negative symptoms **(Figure S7**). Eccentricity scores of thalamic voxels performed moderately when predicting the severity of negative symptoms (r = 0.17 ± 0.25, MAE = 4.47 ± 0.83), while poor in the prediction of positive symptoms (r = 0.06 ± 0.23, MAE = 4.21 ± 0.69). For positive and negative symptoms, predictive models with median performance were separately reported in **Figure 4B**. In the model predicting negative symptoms, weights were heavier in the thalamic voxels regarding to transmodal networks such as the VAN and DMN.

## Discussion

In the current study we investigated macroscale thalamic functional organization in EOS through dimensionality reduction techniques on thalamocortical functional connectivity. We found both expansions along L-M principal axis and A-P secondary axis of thalamic hierarchies in EOS, indicating connectivity profiles between both anchors showed higher dissimilarities in patients versus controls. Disordered functional hierarchies of the thalamus were related to altered thalamocortical interactions both in unimodal and transmodal networks. To evaluate the cytoarchitectural underpinnings of the macroscale functional organization disturbances in EOS, we compared alterations in functional organization within core- and matrix-cells that derived from an independent transcriptomic atlas. We found that in particular, functional organization of thalamic core cells was altered in EOS. Patients’ abnormal coupling patterns between a-priori map of the core-matrix cytoarchitecture and functional connectome characterized a spectrum from perceptual to abstract cognitive functions. Moreover, transcriptomic-informed analyses suggested a close relationship between macroscale functional organization and schizophrenia etiology-related gene expressions in the thalamus. Employing a machine learning strategy, we found that thalamic functional organization was able to predict negative symptoms in EOS. In sum, the current findings provide mechanistic evidence for disrupted thalamocortical system in EOS, and point to alterations in functional networks associated with both perceptual and cognitive functions, suggesting a unitary pathophysiology of heterogeneous symptoms in schizophrenia.

In line with previous work on thalamic hierarchies (25), the principal functional gradient described continuous transition from the ventral lateral nucleus to the anterior and pulvinar groups, and the second axis delineated gradual transition from the anterior nuclei to the pulvinar. Whereas L-M thalamic axis has been reported to correspond to the distribution of gray matter morphology, i.e., low-to-high intensity of neural mass, and the A-P gradient was strongly related to the intrinsic geometry of the thalamus. These findings suggested an association between functional thalamic hierarchies and its structure. Albeit not directly shown, L-M and A-P axes may reflect functional relevance in different dimensions, i.e., two kinds of transitions across functional networks. Indeed, we found that the L-M principal axis functionally segregated the VIS and SMN, similar to the second gradient of cortical connectome (16). Conversely, the A-P axis of thalamic hierarchies segregated unimodal and transmodal networks, in accord with previous findings (25). Taken together, beyond supporting macroscale thalamic hierarchical framework, the current findings further broaden our knowledge of functional specialization of thalamic L-M and A-P axes.

Thalamic ventral lateral and ventral posterior nuclei, as one end of the L-M organizational axis, exhibited evident dissociation in EOS. Both nuclei receive neuronal input from the sensory periphery, and project to the motor and somatosensory cortices, respectively (43). The etiology of schizophrenia has been suggested to damage refinement of motor/somatosensory-thalamic connectivity patterns that occurs during brain maturation (15). Indeed, in adolescent patients relative to adult patients, structural abnormalities in the sensorimotor cortex are reported to be particularly salient (44), but may gradually fade out within a longitudinal period of observation (45). Moreover, motor performance has been reported worse in adolescent patients relative to adult patients when accounting for developmental factors (46). In line with this observation, a meta-analysis suggested that motor deficits may precede the onset of schizophrenia and may constitute robust antecedents of this mental disorder (47). In the context, we postulate sensorimotor-related segregation along thalamic L-M hierarchy might underlie premorbid disturbances in motor development, a marker distinct to schizophrenia.

Compared with TD, EOS patients had increased segregation in two extremes of the A-P axis, i.e., visual-related pulvinar nucleus and default mode-associated anterior nuclear group. Weaker functional connectivity between the VIS and DMN has been previously reported in EOS (48, 49). In fact, given the central role of the thalamus in the development of the cerebral cortex (11), abnormalities of the cerebral cortex in schizophrenia might occur secondary to thalamic pathology (4, 50). Thus, the unimodal-transmodal thalamic hierarchy expansion might further result in disturbed cortical differentiation of unimodal and transmodal regions in EOS. A compression of the unimodal-to-transmodal cortical hierarchy was recently found in chronic adult-onset schizophrenia (51), contrasting with our observation of cortical-thalamic hierarchy expansion. Given age-dependent shifts in the macroscale cortical hierarchy (52), this inconsistence might due to their disparate stage of the illness and age of onset, or their usage of antipsychotic drugs. Further longitudinal works are needed to chart functional organization abnormalities of the thalamus and the cerebral cortex during the course of schizophrenia. Nevertheless, the current findings embed thalamus into a cortical functional organization linked to differentiation of sensory from abstract cognitive functions, paving the way to comprehensively reveal cognitive defects, another well documented precursor of schizophrenia excepts for motor deficits (47).

Leveraging our observations of alterations of thalamic functional organization against a proxy map of core/matrix cells based on post-mortem transcriptomic data (8), we observed thalamic core cells to underlie expansive functional hierarchies in EOS, rather than matrix cells. Core and matrix cells are two primary types of thalamic relay neurons which separately exhibit immunoreactivity to the calcium binding proteins Parvalbumin and Calbindin (9). Thalamic nuclei differ in the ratio of core and matrix neurons (53). Specifically, sensory and motor relay nuclei, as well as the pulvinar nuclei and the intralaminar nuclei are chiefly composed of core cells (54). Compared with matrix cells, the larger core cells innervate middle cortical layers in a more area-restricted and topographically-organized fashion (9). Functionally, Parvalbumin-rich core cells have been reported to act as drivers of feed-forward activity, while Calbindin-rich matrix cells fulfil a more modulatory function (7). Together, this may suggest that thalamic hierarchy disturbances in EOS may relate to the “feed-forward” pathway that transmits information from the sensory periphery, not the “feed-back” pathway (55).

A further inspection of gene-connectome coupling abnormalities in EOS by evaluating the selective connection of the core-matrix cytoarchitecture with the cortex could show that, patients’ core thalamus dys-connected with both unimodal and transmodal cortices. In healthy adults, core regions have preferential connections with unimodal primary regions, and matrix areas with transmodal cortices incorporating the DMN, FPN, VAN and the limbic network (8). Conversely in the current pediatric sample, we observed core-related connections with wide-spread cortical regions including both unimodal and posterior transmodal cortices for TD, whereas EOS had lager similarity with the previously reported adult pattern, i.e., stronger core-related couplings in the VIS, SMN and DAN and weaker couplings in the DMN and FPN. Together, our results imply putatively excessive maturation in the thalamocortical feedforward pathway of schizophrenia. However, future molecular-level work is undoubtedly needed to elaborate on the feedforward pathway alterations in the still developing brain of schizophrenia.

It has been suggested that schizophrenic brain may not form connections according to gene encoded blueprints which have been phylogenetically determined to be the most efficient (56). In line with this, we observed that impaired thalamic hierarchy in EOS was highly associated with schizophrenia-related gene expressions, especially genes encoding Glutamatergic neurotransmission and neurodevelopmental proteins (39). Our findings reveal a gene-connectome correspondence in the thalamocortical system of EOS, adding new evidence for genetics of schizophrenia. However, there is an obvious shortage in our gene-related analyses. The gene expression levels were assessed from post-mortem brain tissue of adults (28), which might be different from the pediatric human brain. Limited by the lack of pediatric transcriptional atlas, our findings about the association between macroscale connectome topology and gene architecture should be carefully considered.

Behavioral decoding of the gene-connectome coupling pattern derived a sensory-cognitive architecture, describing functions associated with primary sensory and multisensory processing to working memory, cognitive control and motivation. Along the continuous behavioral spectrum, perception (especially visual sensory), motor, and higher cognition such as memory are particularly affected by schizophrenia. Consistently, thalamic hierarchy abnormalities were sensorimotor-related, as well as visual/default mode-associated along a second axis. As the “cognitive dysmetria” theory suggested, the multitude and diversity of behavior deficits in schizophrenia might be tied to an impaired fundamental cognitive process mediated by the thalamus (56, 57). This impairment, i.e., cognitive dysmetria referred to a disruption of the fluid and coordinated sequences of both thought and action, leading to a decreased coordination of perception, retention, retrieval, and response functions (58). In particular, our study suggested that abnormal thalamic hierarchy was closely related to negative symptoms of schizophrenia, i.e., a diminution of functions related to motivation and interest. Compared to positive symptoms (such as delusions), negative symptoms are more complex and likely to be the result of systematic disruption. Effective treatment of negative symptoms has long been a clinical challenge for its poor outcomes. The current study provides a thalamic hierarchy framework for heterogeneous behavior deficits related to negative symptoms in schizophrenia, which might denote future therapy of the resistant symptoms.

In sum, the current study describes thalamic functional organization abnormalities in EOS, which could be related to “feed-forward” core thalamocortical pathway. The macroscale disruptions were related to schizophrenia-related genetic factors. Crucially, it might perturb behaviors involving both low-level perception and high-level cognition, resulting in diverse negative syndromes in schizophrenia.

## Data and code availability

The data that support our findings are available from the corresponding author upon reasonable request. The estimated spatial maps of mRNA expression levels were downloaded at: https://www.meduniwien.ac.at/neuroimaging/mRNA.html. The code for functional gradient analysis was adapted from the MICA lab (http://mica-mni.github.io) and is available at https://github.com/Yun-Shuang/Thalamic-functional-gradient-SZ. The code for behavioral decoding was adapted from https://github.com/NeuroanatomyAndConnectivity/gradient_analysis/blob/master/05_metaanalysis_neurosynth.ipynb. Statistical analyses were carried out using PALM (https://fsl.fmrib.ox.ac.uk/fsl/fslwiki/PALM) and BrainSMASH (https://brainsmash.readthedocs.io/). Machine learning analyses were based on scikit-learn package (https://scikit-learn.org/stable/modules/generated/sklearn.linear_model.ElasticNetCV.html). Results were visualized using Connectome Workbench (https://www.humanconnectome.org/software/connectome-workbench), and Seaborn (https://seaborn.pydata.org/) in combination with ColorBrewer (https://github.com/scottclowe/cbrewer2).

## Acknowledgements

We are grateful to all the participants and their guardians in this study. This work was supported by the National Natural Science Foundation of China (82121003, 62036003, 62073058, 62173070), Innovation Team and Talents Cultivation Program of National Administration of Traditional Chinese Medicine (ZYYCXTD-D-202003). S.L.V. was also funded in part by Helmholtz Association’s Initiative and Networking Fund under the Helmholtz International Lab grant agreement InterLabs-0015, and the Canada First Research Excellence Fund (CFREF Competition 2, 2015-2016) awarded to the Healthy Brains, Healthy Lives initiative at McGill University, through the Helmholtz International BigBrain Analytics and Learning Laboratory (HIBALL).

