## supplement for "Macroscale Thalamic Functional Organization Disturbances and Underlying Core Cytoarchitecture in Early-Onset Schizophrenia"

Running title: Expansion of thalamic functional hierarchies in schizophrenia

Yun-Shuang Fan ^1,2^, Yong Xu ^8^, Şeyma Bayrak ^2^, James M. Shine ^3^, Bin Wan ^2,5,7^, Haoru Li ^1^, Liang Li ^1,6^, Siqi Yang ^1^, Yao Meng ^1^, Sofie Louise Valk ^2,5*#^, Huafu Chen ^1,4*#^

*1.* *The Clinical Hospital of Chengdu Brain Science Institute, School of Life Science and Technology, University of Electronic Science and Technology of China, Chengdu, China; 2. Otto Hahn Group Cognitive Neurogenetics,* *Max Planck Institute for Human Cognitive and Brain Sciences, Leipzig, Germany; 3. Brain and Mind Center, The University of Sydney, Sydney, Australia; 4. MOE Key Lab for Neuroinformation, High-Field Magnetic Resonance Brain Imaging Key Laboratory of Sichuan Province,* *University of Electronic Science and Technology of China, Chengdu, China; 5. Institute of Neuroscience and Medicine (INM-7: Brain and Behavior), Research Centre Jülich, Jülich, Germany; 6. Academy for Advanced Interdisciplinary Studies, Peking University; 7. International Max Planck Research School on Neuroscience of Communication: Function, Structure, and Plasticity (IMPRS NeuroCom), Leipzig, Germany; 8. Department of Psychiatry, First Hospital/First Clinical Medical College of Shanxi Medical University, Taiyuan, China.*

* Both last co-authors contributed equally.

\# Corresponding authors:

Huafu Chen, & Sofie Louise Valk,.

**Supplement 1**

**MRI preprocessing**

T1-weighted structural data were preprocessed with FreeSurfer (v7.1.0, http://surfer.nmr.mgh.harvard.edu/), which included cortical segmentation and surface reconstruction. Rs-fMRI functional data were preprocessed with the CBIG pipeline (https://github.com/ThomasYeoLab/CBIG) based on FSL [v5.0.9, (1)] and FreeSurfer (v7.1.0), which included removal of the first four volumes, slice-timing, motion correction, and boundary-based registration to structural images. Participants with the intrasubject registration cost exceeding the 0.7 were discarded. Additionally, subjects with severe segmentation faults were excluded via visual inspection. Volumes with FD > 0.2 mm or voxel-wise differentiated signal variance > 50 were marked as outliers. Censored images included one volume before and two volumes after outliers, as well as images lasting fewer than five contiguous volumes time frames. Participants with a mean FD > 0.2 mm or censored volumes percent > 50% were excluded from all further analyses. Censored images were excluded and substituted by the least square interpolation of neighbor time frames. White matter signal, ventricular signal, head motion parameters and temporal derivatives were regressed out to remove spurious noise affects. Subsequently, functional images were then bandpass filtered (0.01–0.08 Hz). Preprocessed images were then registered to MNI152 template and resampled to the cortical surface using Ciftify package [v2.3.3, (2)]. The thalamus was localized using the Gordon 333 Atlas (3), including 2536 voxels across both hemispheres.

**Macroscale thalamocortical gradients identification**

Analogous to previous work (4), gradients of thalamocortical functional connectome were generated using the diffusion embedding algorithm in BrainSpace Toolbox (5). First, thalamocortical functional connectome was calculated based on Pearson correlations between the thalamic and cortical rs-fMRI time-series for each subject (6, 7). A group representative connectome was calculated by averaging thalamocortical connectomes across patients and controls. Both individual-level and group-level connectome matrices were then row-wise thresholded (the top 10% connections retained) and converted into cosine similarity matrices (6, 8). Subsequently, nonlinear dimensionality reduction techniques were employed on the group-level similarity matrix to resolve a template connectome gradient, i.e., spatial axis in connectome variations (9). Individual gradients were then estimated and aligned to the template gradients. To evaluate the influence of the alignment template, we conducted a replication by using mean thalamocortical connectome only across controls as the group-level template in gradient analysis. Replication results indicated that our main findings were stable and unaffected by the alignment template (**Figure S1**). Finally, they were smoothed (FWHM = 5mm) across thalamic voxels to avoid introducing artifactual correlation. We selected the first two gradients to represent the macroscale thalamic connectome manifolds for their individual eigenvariance > 10%, which explained 44% of the total eigenvariance in functional connectome (**Figure S2**).

**The core-matrix cytoarchitecture**

***Linking thalamic core-matrix cytoarchitecture with connectome manifolds***

To delineate the core-matrix cytoarchitecture in the thalamus, we used the spatial maps of mRNA expression levels for two calcium-binding proteins (CALB1 and PVALB) (https://github.com/macshine/corematrix) generated from post-mortem Allen Human Brain Atlas (10). Two (CUST_11451_PI416261804 and A_23_P17844) and three probes (CUST_140_PI416408490, CUST_16773_PI416261804 and A_23_P43197) were used to estimate the gene expressions of PVALB and CALB1, respectively (11). Expression differences between normalized CALB1 and PVALB levels denoted CP index, i.e., CALB1-PVALB values for thalamic voxels. Thalamic voxels with positive CP values related to matrix projection cells, and voxels with negative values related to core populations. Global eccentricity indices of core/matrix populations were separately measured by averaging eccentricity values across all thalamic core/matrix cells, and compared with each other.

Subsequently, we sought to reveal an association between the thalamic cytoarchitecture and disturbed connectome manifold in early-onset schizophrenia (EOS) patients. Between-group difference map of eccentricity was used as quantification index for perturbances of macroscale connectome gradients. Spatial correlations were employed between gene expression maps and differential eccentricity map between the EOS and typically developing (TD) groups. To correct for the spatial autocorrelation (SA) of brain maps, we employed the variogram-matching model to simulated surrogate maps (N = 1,000) preserving SA of empirical volumetric maps (12). These surrogate maps were then used as null distributions for evaluating SA-corrected statistical significance.

***Behavioral decoding of gene-connectome couplings***

To estimate behavioral implications of couplings between core-matrix cytoarchitecture and functional connectome, we projected CP maps onto the cerebral cortex. For each subject, gene-connectome coupling map was computed by spatially correlating CP map and thalamocortical connectome (11), and were then down sampled into 400 cortical parcels according to the Schaefer atlas (13). Cortical parcels with positive coupling values indicated as preferential associations with matrix thalamic populations, and negative values suggested core populations.

Next, topic-based behavioral decoding was conducted to evaluate cognitive terms associated with coupling maps using NeuroSynth meta-analytic database (14). The mean coupling map across all subjects was divided into five-percentile bins and binarized for creating regions of interest (ROIs) masks in the meta-analyses. Topics were derived from the 50-set topic, where 6 topics were removed as “noise” according to a previous study (6). Twenty-one cognitive topics remained after thresholding of z−statistic > 3.1. Core populations preferentially linked with unimodal primary regions that subserve primary sensory and multisensory functions, and matrix areas to transmodal cortices characterized by more abstract cognition (**Figure S6**), suggesting a sensory-cognitive behavioral architecture. Along the continuous spectrum, cognitive consequences shifted from primary sensory and multisensory processing to working memory, cognitive control and motivation.

Moreover, gene-connectome coupling map with significant group differences between patients and controls was binarized for creating regions of interest (ROIs) masks in the meta-analyses. This analysis was constrained in the 21 cognitive topics significantly associated with gene-connectome coupling maps (z−statistic > 3.1). To better visualize our result, we generated a word cloud (https://amueller.github.io/word_cloud/) summarizing frequency of all cognitive terms included in associated topics as previously suggested (15). The size of a cognitive term in the word cloud is proportional to the strength of its loading.

**Clinical symptoms prediction**

To investigate clinical correspondences of thalamic functional organization, we further used the elastic net model, i.e., a linear regression model that incorporates both Lasso-regularization and Ridge-regularization, to predict clinical symptoms in EOS. The algorithm was implemented in Python using the scikit-learn package (16). Eccentricity values of the two-dimensional gradient space were defined as input features, and PANSS positive and negative scores were used as predictors. A total of 65 patients were randomly split into training (80%) and testing (20%) datasets. The model parameters were optimized through a 5-fold cross-validation. Briefly, the training set was equally separated into five parts: four for the model estimation, one for the validation, which were repeated five times to traverse all parts for validation. The model with the highest cross-validation accuracy was then applied to the testing set to estimate their clinical scores. The model performance was evaluated by comparing observed and predicted clinical scores using mean absolute error (MAE). This procedure including model learning and testing was repeated 101 times, resulting 101 fitted models. The model with median performance was reported.

**Supplement 2**

**Cortical-thalamic gradient identification**

To compare with previous cortical gradient findings in schizophrenia, we also calculated cortical-thalamic gradients based on our thalamocortical functional connectomes. Averaged thalamocortical connectome across patients and controls was also used as group representative connectome. For each cortical vertex, both individual-level and group-level connectome matrices were then column-wise thresholded (the top 10% connections retained) and converted into cosine similarity matrices. Nonlinear dimensionality reduction techniques were employed on cortical similarity matrices to generate cortical-thalamic gradient as our thalamocortical gradient identification.

**Cortical-thalamic gradient findings**

From a visual perspective, the first cortical gradient (28% eigenvariance) captured a unimodal-to-transmodal axis of cortical-thalamic connectivity, where the default mode network (DMN) was situated on one side (**Figure S3A**). Compared with TD controls, the first gradient axis of EOS patients was extended, especially at the end regarding to the DMN. Unexpectedly, the second cortical gradient (14% eigenvariance) ran along an anterior-to-posterior axis, one end of which was anchored at the ventral attention network (VAN) (**Figure S3B**). In line with our thalamo-cotrical gradient results, patients had an expansion of this macroscale cortical functional organization, indicating greater functional segregation compared to controls (**Figure S3C**).

**Tables**

**Table S1.** 28 relevant protein coding genes for schizophrenia (17).

|  | **Name** | **location** | **Thalamic**  **Expressions** | **Functions** |
| --- | --- | --- | --- | --- |
| Therapeutic targets | | | | |
| 1 | DRD2 | 11q23.2 | 0.14±0.03 | Blockade of the dopamine type 2 receptor subtype is a necessary and sufficient condition for antipsychotic activity. |
| 2 | GRM3 | 7q21.12 | 0.35±0.11 | mGluR3 is a metabotropic glutamate receptor treated as a potential therapeutic target. |
| Glutamatergic neurotransmission | | | | |
| 3 | GRIN2A | 16p13.2 | 0.28±0.09 | The NMDA receptor subunit GRIN2A (NR2A) is a key mediator of synaptic plasticity. |
| 4 | GRIA1 | 5q33.2 | 0.24±0.06 | Glutamate receptor 1 (GluR1, GluA1) is a subunit of an AMPA (non-NMDA) receptor that mediates fast synaptic transmission. |
| 5 | SRR | 17p13.3 | 0.16±0.03 | Serine racemase catalyzes L-serine racemization to D-serine, an essential coagonist and activator of NMDA receptors. |
| 6 | CLCN3 | 4q33 | 0.13±0.02 | CLC-3 is a voltage-gated chloride channel localized to glutamatergic synapses in the hippocampus, where it modulates plasticity. |
| Neuronal calcium signaling | | | | |
| 7 | CACNA1I | 22q13.1 | 0.10±0.01 | When CACNA1I co-activates with NR2B-containing NMDA receptors, activation triggers synaptic plasticity and long-term potentiation. |
| 8 | RIMS1 | 6q12-13 | 0.29±0.08 | RIMs are multi-domain proteins that tether calcium channels to synaptic active zones, dock and prime synaptic vesicles for release, mediate presynaptic plasticity and facilitate neurotransmitter release. |
| Synaptic function and plasticity | | | | |
| 9 | KCTD13 | 16p11.2 | 0.15±0.03 | Polymerase Delta-Interacting Protein 1 lies within a pathogenic CNV at 16p11.2 associated with neurodevelopmental disorders and brain and body size phenotypes. |
| 10 | NLGN4X | Xp21.33-32 | 0.18±0.03 | Nlgn4 may modulate the pre-synaptic calcium channel population through its interaction with neurexins. |
| 11 | IGSF9 | 11q25 | 0.09±0.01 | IgSF9b is strongly expressed in GABAergic interneurons, localized to hippocampal and cortical inhibitory synapses. |
| 12 | CNTN4 | 3p26.3 | 0.36±0.12 | Contactins are axon-associated cell adhesion molecules that function in neuronal network formation and plasticity. |
| 13 | MEF2C | 5q14.3 | 0.28±0.08 | It is a transcription factor regulating neurogenesis, excitatory synapse number, dendrite morphogenesis and differentiation of post-synaptic structures. |
| 14 | PTN | 7q33 | 0.08±0.01 | Pleiotrophin is a developmentally regulated neurite growth-promoting factor (NEGF) family cytokine/growth factor. |
| 15 | CNKSR2 | Xp22.12 | 0.23±0.06 | CNK2 plays a role in assembly of synaptic complexes at the postsynaptic membrane and coupling of signal transduction to membrane/cytoskeletal remodelling. |
| 16 | PAK6 | 15q14 | 0.28±0.08 | PAK6 is a highly brain expressed serine/threonine protein kinase associated  with neurite outgrowth, filipodia formation and cell survival. |
| 17 | SNAP91 | 6q14.2 | 0.19±0.04 | Together with CALM, SNAP91 (AP180) establishes the polarity and controls the growth of axons and dendrites in embryonic hippocampal neurons. |
| Other neuronal ion channels | | | | |
| 18 | KCNB1 | 20q13.13 | 0.26±0.08 | Kv2.1 is abundantly expressed in the cortex and hippocampus, where it regulates neuronal excitability, action potential duration, and tonic spiking. |
| 19 | HCN1 | 5p21 | 0.21±0.04 | HCN1 is a major contributor to the inward hyperpolarization-activated cation current (Ih) current in the brain, which regulates neuronal excitability, rhythmic activity and synaptic plasticity. |
| 20 | CHRNA3 | 15q25.1 | 0.46±0.18 | It is a nicotinic acetylcholine receptor (nAChR) forming ligand-gated ion channels in certain neurons and also on the presynaptic and postsynaptic sides of the neuromuscular junction. |
| 21 | CHRNA5 | 15q25.1 | 0.24±0.05 | CHRNA5 is also a type of nAChR genes. |
| Neurodevelopment | | | | |
| 22 | FXR1 | 3q26.33 | 0.20±0.04 | FXR1P is found in dendritic spines in the mouse hippocampus and targets mRNAs and microRNAs including brain specific miRNA9 and miR-24. |
| 23 | SATB2 | 2q33.1 | 0.18±0.05 | SATB2 is a DNA binding protein that binds nuclear matrix attachment regions, regulating transcription and chromatin remodeling. |
| 24 | PODXL | — | 0.12±0.02 | — |
| 25 | BCL11B | — | 0.50±0.20 | — |
| 26 | TLE1 | — | 0.14±0.02 | — |
| 27 | TLE3 | — | 0.18±0.02 | — |
| 28 | FAM5B | — | 0.29±0.07 | — |

Note: Mean ± SD.

**Figures**

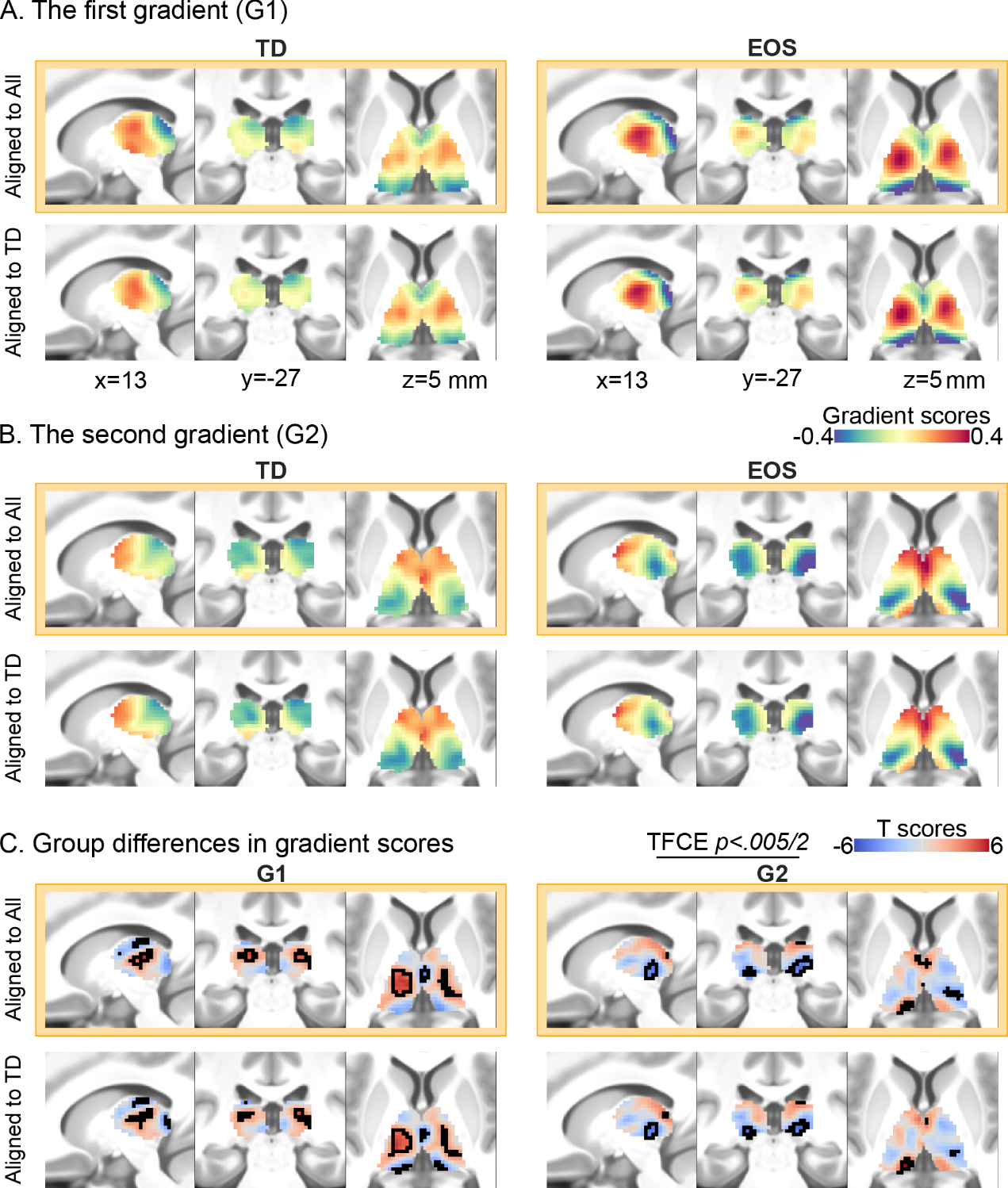

**Figure S1. Replication of main results.** To exclude the influence of alignment template in gradient analysis, we also used mean TD thalamocortical connectome to calculate the template gradients. These results replicated our main results (outlined in orange), thereby indicating that these findings were stable and unaffected by the alignment template. TD, typically developing; EOS, early-onset schizophrenia.

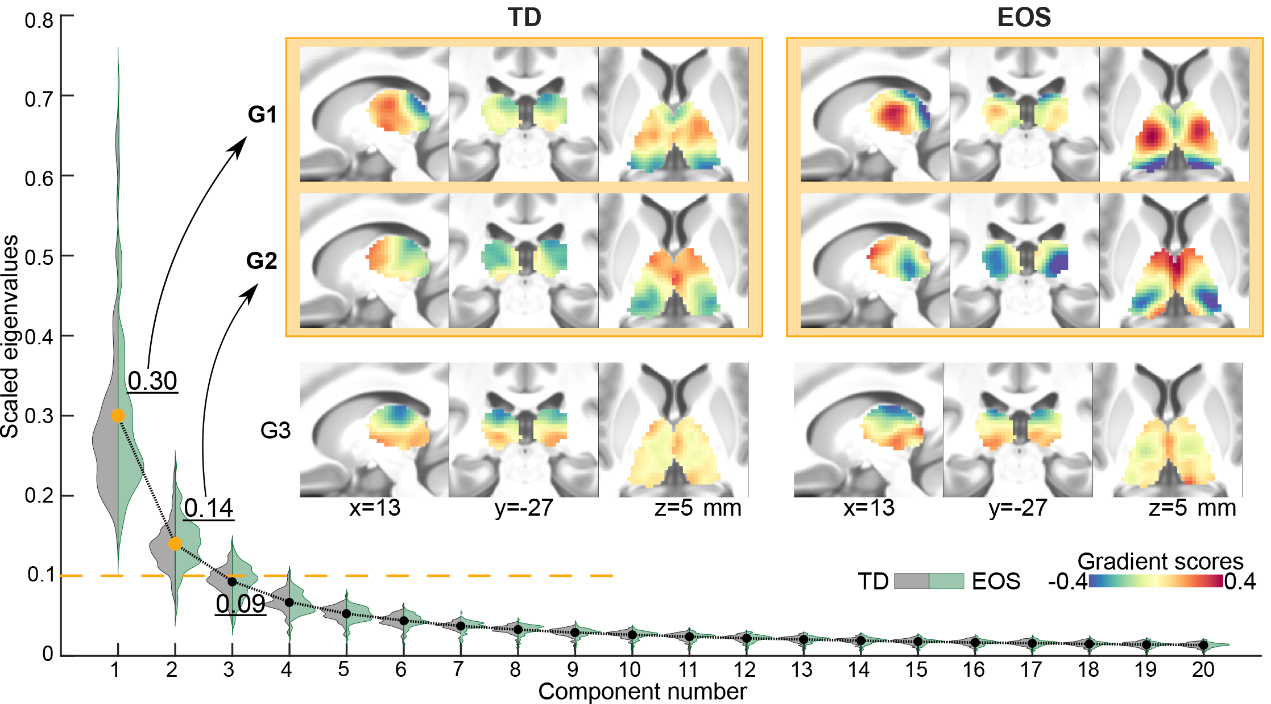

**Figure S2. Eigenvariance explained by the first three gradients.** The first (30%) and the second gradients (14%) were selected out for their eigenvariances larger than 10%. Additionally, the third gradients in the TD and EOS groups were shown below the first two gradients, which tends to be a dorsal-ventral thalamic axis.

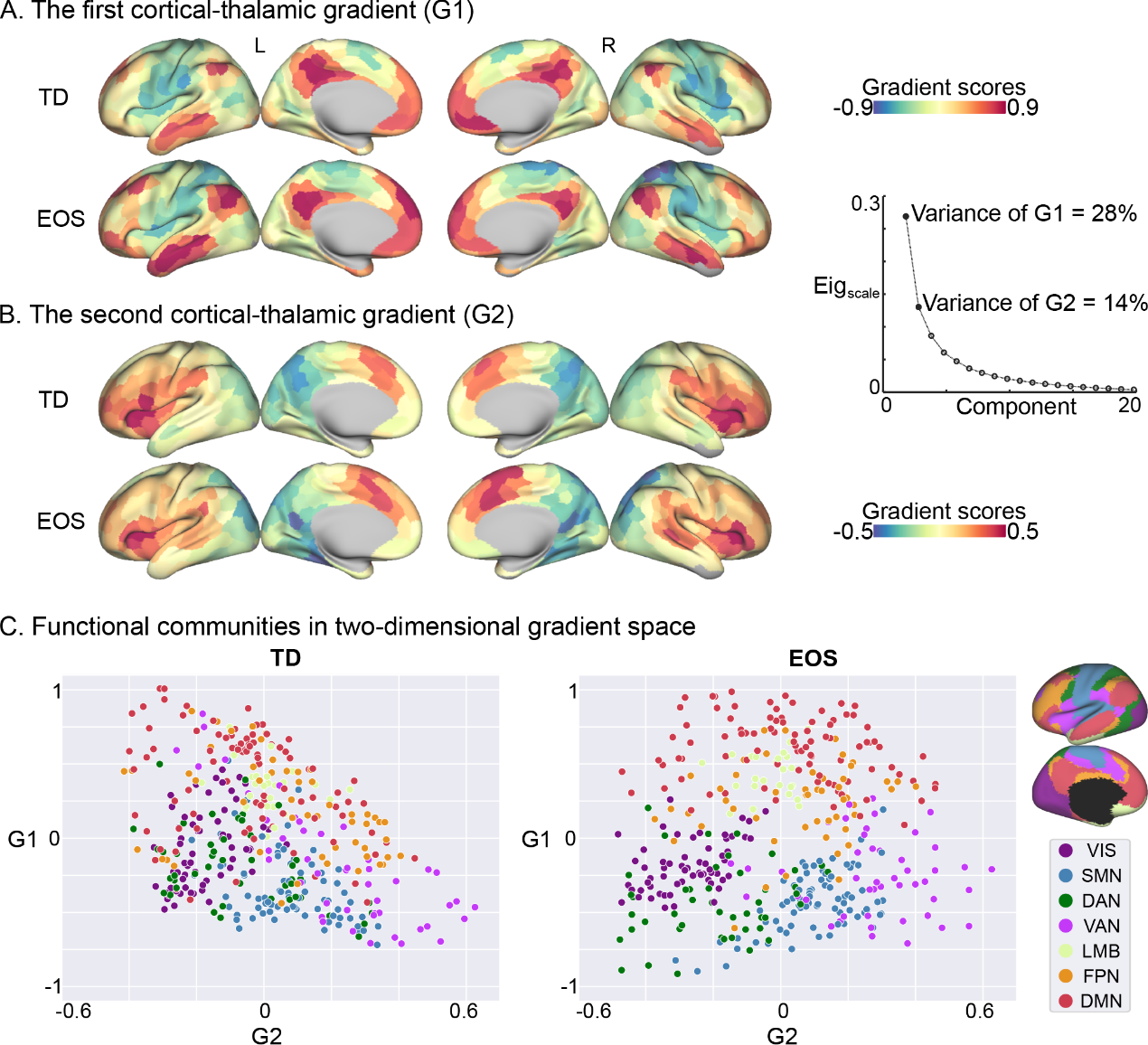

**Figure S3. Cortical-thalamic gradients in two groups.** Cortical G1 and G2 explains 28% eigenvariance and 14% separately. G1 differentializes unimodal (including VIS, SMN, DAN, VAN) with transmodal networks (including DMN, FPN, LMB). The anterior-to-posterior G2 is more likely to isolate VAN from other networks. In the two-dimensional gradient, EOS patients’ DAN is visually more expanded relative to TD controls.

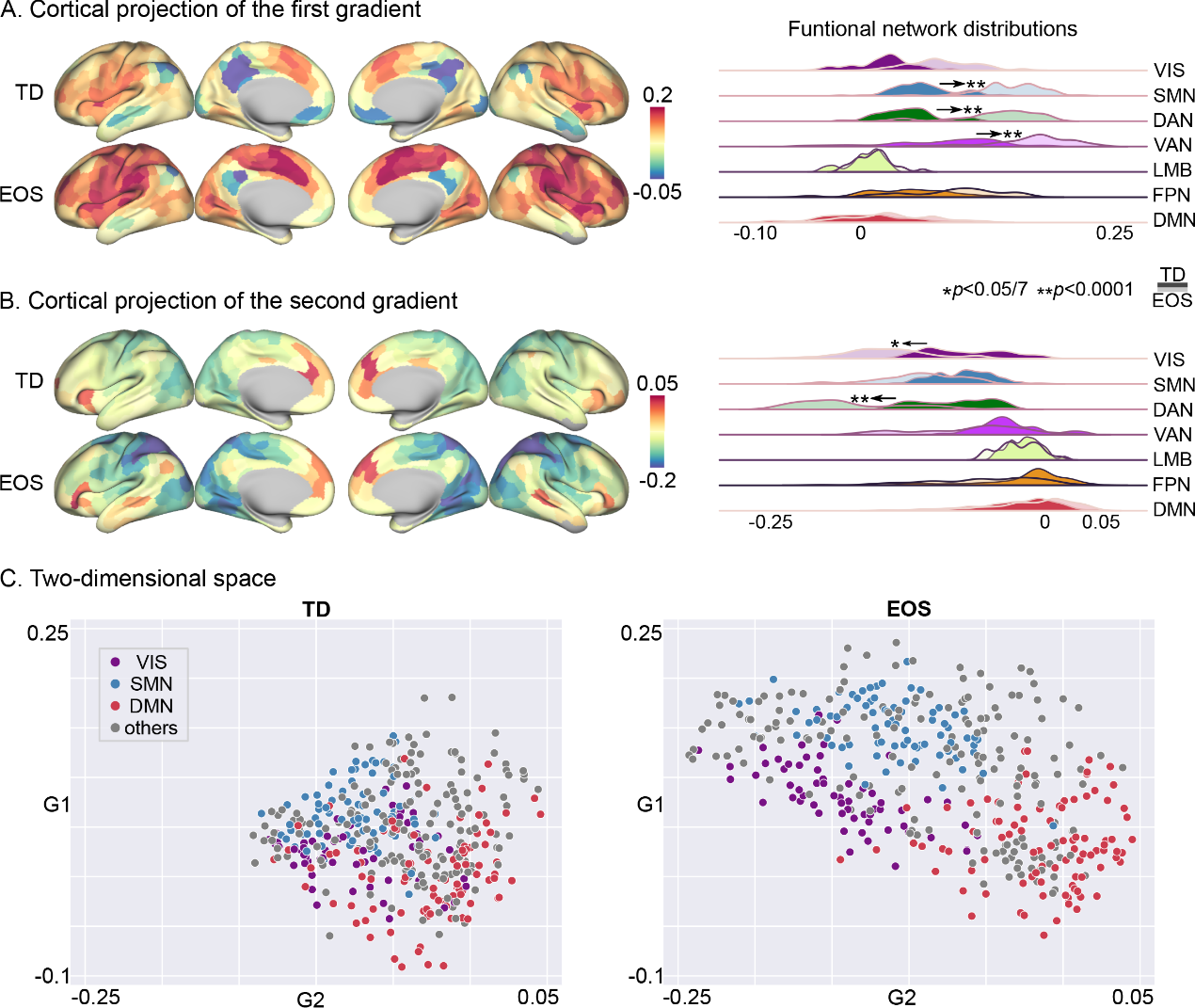

**Figure S4. Projecting thalamic gradients to the cerebral cortex.** Density maps indicate mean gradient scores within cortical functional networks for EOS (lighter) and TD (darker). Network level differences between EOS and TD were accessed by t-tests, and significant differences were depicted by * (Bonferroni correction, *p* < 0.05) and ** (*p* < 0.0001). VIS, the visual network; SMN, the sensorimotor network; DAN, the dorsal attention network; VAN, the ventral attention network; LMB, limbic network; FPN, the frontoparietal network; DMN, the default mode network.

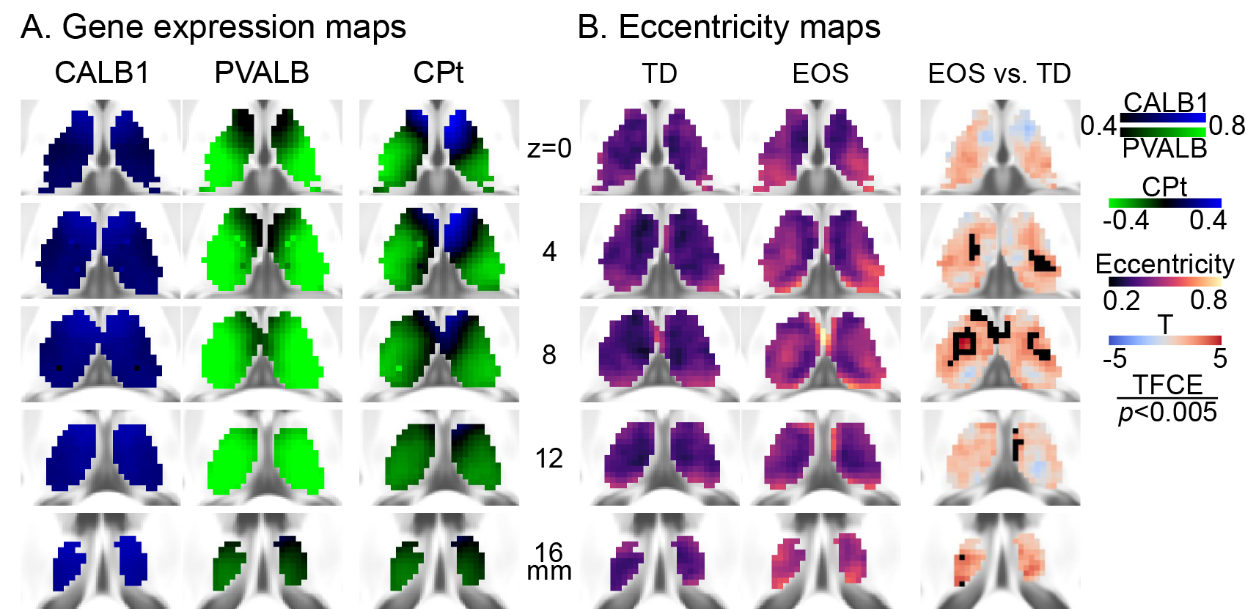

**Figure S5. The core-matrix cytoarchitectural and thalamic gradient eccentricity maps. (A)** mRNA expression levels for Calbindin (CALB1) and Parvalbumin (PVALB) proteins, and eccentricity scores for thalamic voxels. CP denotes the differential gene expression level for CALB1 and PVALB. Thalamic voxels with positive CP values (blue) indicate matrix projection cells, and negative CP values (green) related to core populations. **(B)** Eccentricity scores of thalamic voxels are depicted for TD and EOS, and their differences are assessed by t-tests (EOS vs. TD). Voxels with significant eccentricity differences are surrounded by black contours (TFCE, *p* < 0.005).

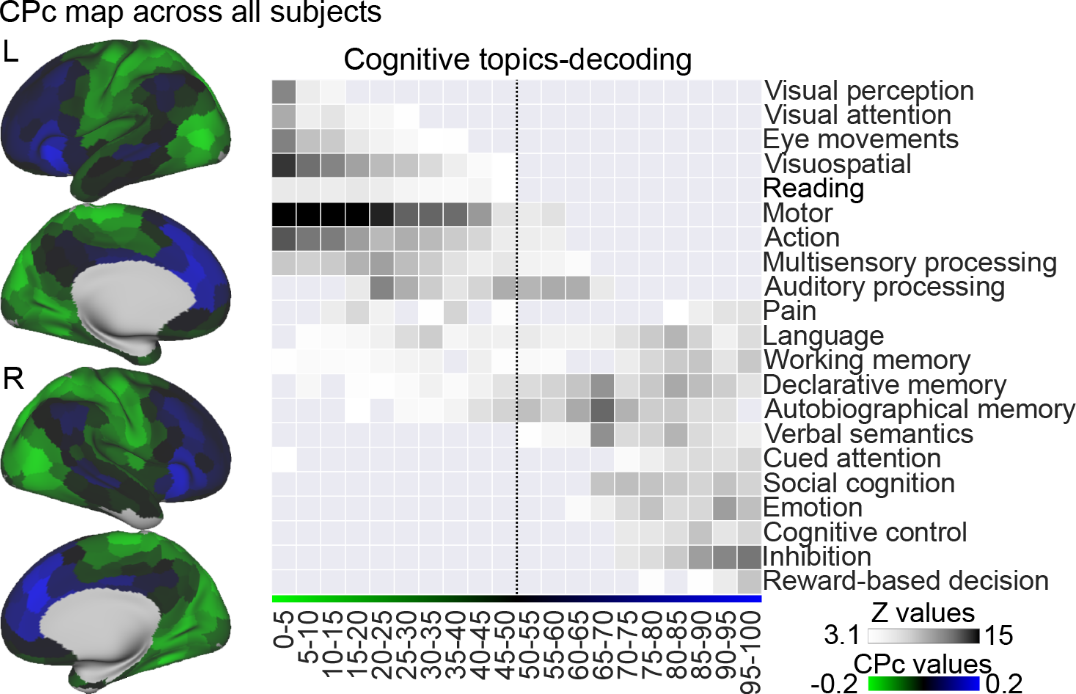

**Figure S6. Cognitive consequences of gene-connectome coupling.** Parcel-wise coupling map (left) was obtained using all subjects including patients and controls. In the right panel, each cell is colored according to their z-statistic values. Each row represents a cognitive topic, and each column represents a five-percentile bin of coupling mask. Dotted line depicts where coupling value is equal to zero.

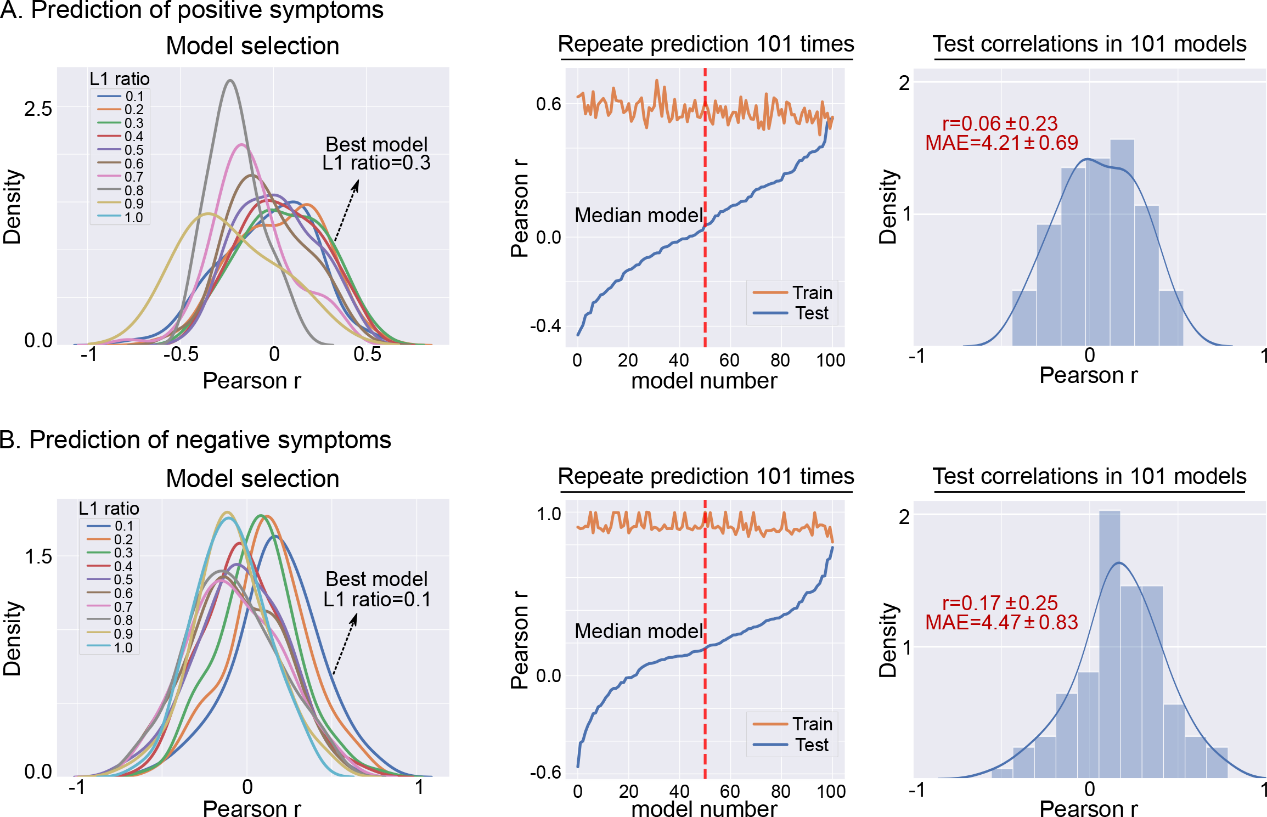
**Figure S7. Predictive model selection. (A)** For predicting PANSS positive scores, the regression model with L1 ratio equal to 0.3 was the best model (left). This prediction procedure was repeated 101 times, resulting 101 models, where the model with median performance was reported (middle). The Pearson correlation between observed and predicted clinical scores was used to evaluate the model performance (right). **(B)** The regression model with L1 ratio equal to 0.1 was the best model to predict PANSS negative scores. MAE, mean absolute error.
